# Membrane bending begins at any stage of clathrin-coat assembly and defines endocytic dynamics

**DOI:** 10.1101/163303

**Authors:** Brandon L. Scott, Kem A. Sochacki, Shalini T. Low-Nam, Elizabeth M. Bailey, QuocAhn Luu, Amy Hor, Andrea M. Dickey, Steve Smith, Jason G. Kerkvliet, Justin W. Taraska, Adam D. Hoppe

## Abstract

Clathrin-mediated endocytosis internalizes membrane from the cell surface by reshaping flat regions of membrane into spherical vesicles(1, 2). The relationship between membrane bending and clathrin coatomer assembly has been inferred from electron microscopy and structural biology, without directly visualization of membrane bending dynamics (3–6). This has resulted in two distinct and opposing models for how clathrin bends membrane (7–10). Here, polarized Total Internal Reflection Fluorescence microscopy was improved and combined with electron microscopy, atomic force microscopy, and super-resolution imaging to measure membrane bending during endogenous clathrin and dynamin assembly in living cells. Surprisingly, and not predicted by either model, the timing of membrane bending was variable relative to clathrin assembly. Approximately half of the time, membrane bending occurs at the start of clathrin assembly, in the other half, the onset of membrane bending lags clathrin arrival, and occasionally completely assembled flat clathrin transitions into a pit. Importantly, once the membrane bends, the process proceeds to scission with similar timing. We conclude that the pathway of coatomer formation is versatile and can bend the membrane during or after the assembly of the clathrin lattice. These results highlight the heterogeneity in this fundamental biological process, and provide a more complete nanoscale view of membrane bending dynamics during endocytosis.

---

Early observations by electron microscopy suggested that clathrin oligomerizes into flat hexagonal lattices on the plasma membrane that rearrange into spheres via the transition of some of the hexagons into pentagons(9). However, structural biology arguments suggest that this conversion would be energetically costly; leading to a model in which membrane bending occurs progressively with the assembly of clathrin(7, 8) (Fig. 1a). Recently, thin-section correlative fluorescence-cryoelectron microscopy support the model of pre-assembled clathrin sheets that subsequently bend into vesicles(10) (Fig. 1b). Thus, there remains uncertainty in the fundamental mechanism by which clathrin-coat assembly bends membrane in mammalian cells.

**Fig. 1.**
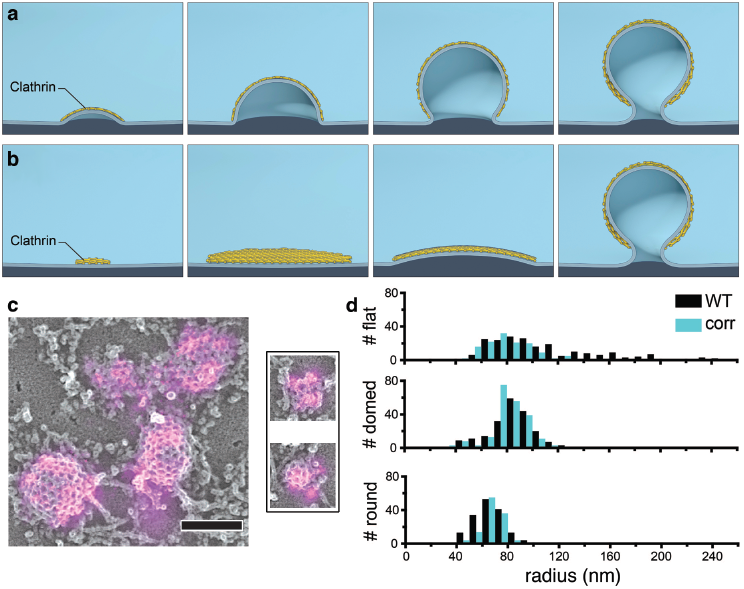
Mechanisms of membrane bending during CME and Clathrin ultrastructure in unroofed SK-MEL-2 cells. **a.** Schematic representation of CME in which membrane bending proceeds with a fixed radius of curvature during the addition of clathrin subunits. **b.**Representation of CME in which clathrin first assembles into a flat sheet that remodels into a vesicle. **c.** Correlative dSTORM and platinum-replica TEM images of fluorescently-labeled clathrin (magenta) demonstrate a range of heterogeneous topographies even at the earliest stages of CME; scale bar is 200 nm. **d.** The size distribution observed amongst clathrin structures in WT SK-MEL-2 cells (black) or SK-MEL-2 cells exogenously expressing clathrin light chain for correlative microscopy (blue). The inset shows ratios of average fluorescence associated with round vs. domed structures and domed vs. flat structures (SD shown for N=3 cell membranes).

The two membrane bending models exhibit distinct and opposing relationships between the changing pit size and clathrin assembly. We applied single molecule super-resolution immunofluorescence of clathrin light chain combined with platinum-replica correlative electron microscopy to image the structure of even the smallest clathrin-coated assemblies at the plasma membrane of SK-MEL-2 cells (Fig. 1c). The fluorescence signal increased over flat to domed to highly invaginated clathrin structures indicating that clathrin was added throughout pit formation (Fig. 1d). Additionally, the observed clathrin morphologies were heterogeneous and displayed a range of lateral radii (Fig. 1d, and Extended Data Figures 3,4), raising the possibility that clathrin accommodates multiple modes of membrane bending and the addition of new clathrin subunits at different stages(11, 12). Measuring the dynamics of membrane bending during clathrin assembly at single endocytic sites in living cells is required to distinguish the possible modes of membrane bending.

Polarized Total Internal Reflection Fluorescence (pol-TIRF) microscopy generates contrast between vertical and horizontal DiI-C18 labeled plasma membrane in living cells(13–15). Pol-TIRF has been used to image changes in membrane topography during exocytosis of chromaffin granules, which are much larger than CME sites (16–18). We improved upon these methods by developing a microscope capable of creating pol-TIRF fields that were parallel (s-pol, S) or perpendicular (p-pol, P) to the coverslip with improved spatial uniformity by averaging multiple illumination directions(19, 20) (Extended Data Figure 2). These P and S fields were used to selectively excite DiI molecules in vertical or horizontal membrane respectively, thereby encoding membrane topography into the ratio of P/S fluorescence images from DiI (Fig. 2a). A computer simulation of pol-TIRF for the formation of 100 nm vesicles by either model (Fig. 1a,b) predicted that the P/S image was sensitive to small changes in membrane bending (Fig. 2c).Although the simulation predicted small differences between the two CME models, these models can be readily distinguished by comparing P/S with the arrival of clathrin (Fig. 2b, and Extended Data Figure 1). For membrane bending during assembly, clathrin and P/S increase together as the pit forms (Fig. 2b). Conversely, for the model in which flat clathrin is reshaped into a sphere, the clathrin intensity is maximal prior to changes in P/S and then decreases as bending moves the top of the pit deeper in the exponentially decaying TIRF field (Fig. 2b, and Extended Data Figure 1). We considered the possibility that detection of the P/S signal would be less sensitive than detection of fluorescent clathrin arrival, thereby creating an apparent temporal delay between clathrin arrival and membrane bending. However, simulations of P/S and clathrin over a range of signal-to-noise ratios (SNR) indicated that detection of P/S is weakly dependent on the SNR and that the two models are distinguishable over a wide range of SNRs encountered in live-cell imaging (Fig. 2b, and Extended Data Figure 1).

**Fig. 2.**
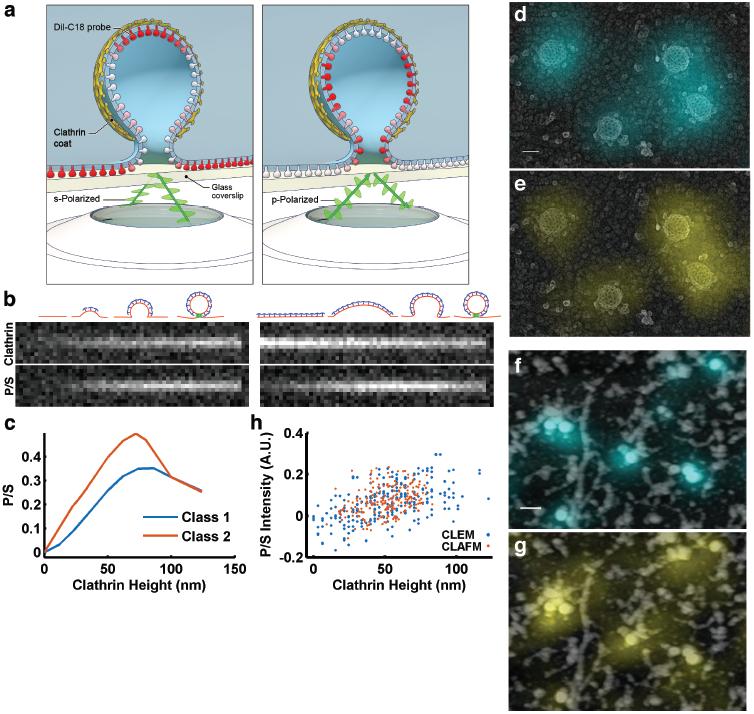
Polarized TIRF microscopy enables imaging of membrane bending at clathrin coated structures. **a.** Schematic representation of pol-TIRF. DiI–C18 orients its dipole moment with the plasma membrane. S-polarized TIRF illuminates horizontal dye molecules, whereas P-polarized TIRF selectively excites vertical dye molecules. The P/S provides contrast for membrane bending. **b.** Simulation of pol-TIRF signals for Class 1 and Class 2 membrane bending. **c**. Quantification of high-resolution simulation at 10 discrete points for Class 1 (blue) and Class 2 (orange) in the absence of noise. **d.** Correlative TEM-pol-TIRF imaging. Overlay of fluorescence from endogenous clathrin-Tq2 on the TEM micrograph showing four clathrin structures; scale bar is 100 nm. **e.** Overlay of P/S on the same region of the micrograph. **f**. Correlative AFM-pol-TIRF imaging; Overlay of clathrin-Tq2 on AFM micrograph; scale bar is 250 nm. **g.** Overlay of P/S signal on the same region of the AFM micrograph. **h.** Quantification of P/S intensity and heights from correlative tomographic reconstructions (blue), and AFM (orange).

We validated pol-TIRF’s sensitivity to membrane bending during CME by correlative light and electron (CLEM) and light and atomic force microscopy (CLAFM). In pol-TIRF-CLEM, endogenous clathrin-Tq2 fluorescence colocalized with the expected clathrin ultrastructure (Fig. 2d), and corresponding P/S signals were observed on individual clathrin-coated pits over a range of invagination stages (Fig. 2e). Importantly, P/S increased with pit stage as determined by morphology (Extended Data Figure 3), and with pit heights determined from TEM tomograms (Fig. 2h, ρ = 0.548, p<0.001, and Extended Data Figure 4). Pol-TIRF-CLAFM on wet samples confirmed pol-TIRF’s sensitivity for membrane bending, despite the reduced resolution achieved by AFM owing to the softness of biological membranes.Specifically, the endogenous clathrin-Tq2 and P/S overlaid with peaks in the AFM images (Fig. 2f, g), and pit height and P/S were positively correlated (Fig. 2h, ρ = 0.351, p<0.001, and Extended Data Figure 4). Variations in CLEM and CLAFM measurements and topographical features near the measured pit were responsible for variability in P/S ratio quantification (Fig. 2h,and Extended Data Figure 4). Therefore, pol-TIRF is sensitive to membrane bending on the scale of CME.

**Fig. 3.**
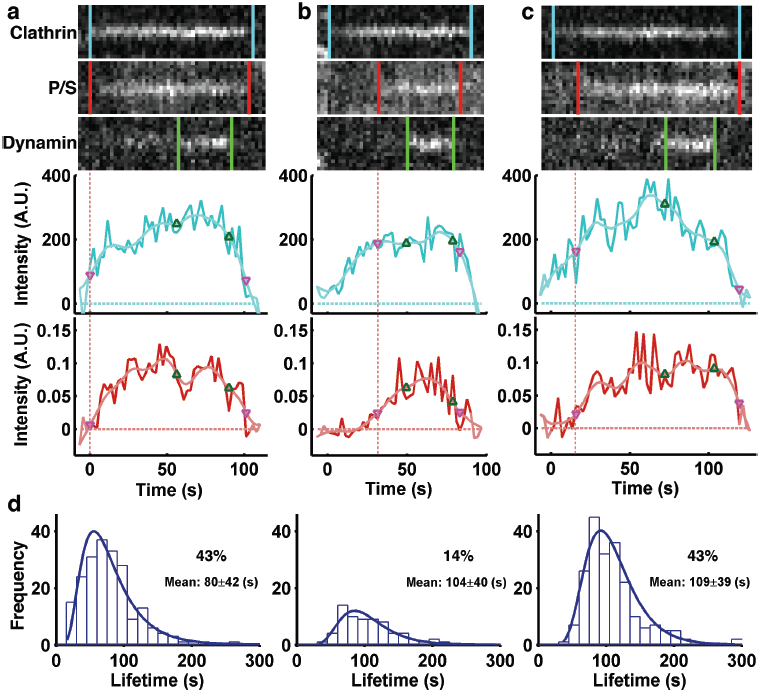
Distinct modes of membrane bending observed by pol-TIRF. **a-c.** Kymographs of clathrin-Tq2, membrane bending (P/S), dynamin-eGFP, and corresponding intensity traces for clathrin (cyan) and P/S (red) for the three classes of membrane bending observed by pol-TIRF. The colored lines indicate the lifetime for each event, the dashed red line highlights the start of P/S event. **a.** Class 1: Clathrin and P/S signals proceed together indicating membrane bending as clathrin assembles. **b.** Class 2: Clathrin signal plateaus prior to the start of P/S indicating all required clathrin was present as a flat sheet. **c.** Class 3: Clathrin assembles prior to P/S signal, but new clathrin was recruited as the membrane bends and the vesicle is formed. **d.** The clathrin lifetime histogram of each class from an average of 4 cells (N = 481 tracks total).

**Fig. 4.**
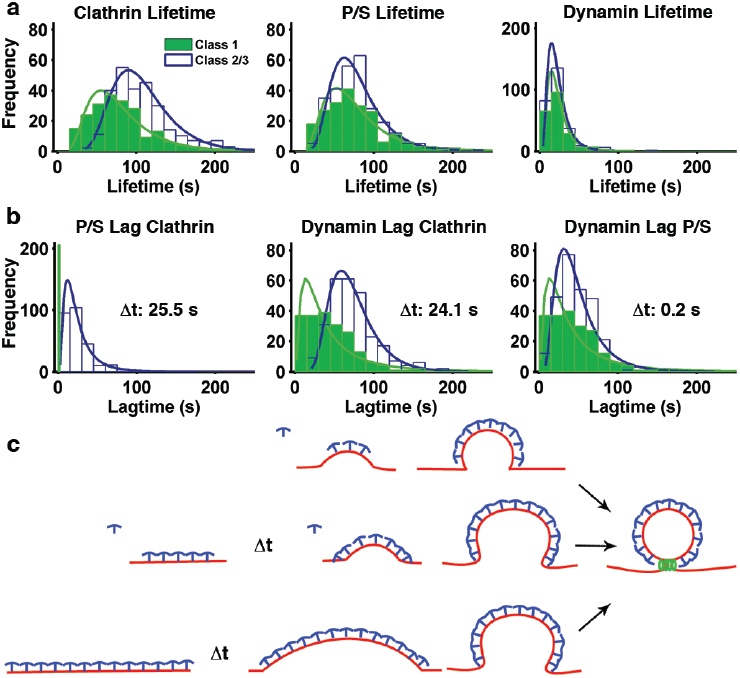
Lifetime analysis of clathrin events. **a.** Lifetime distribution for clathrin, P/S, and dynamin; Class 1, green bars and line, Class 2/3 open with blue line. **b.** Lag time for P/S and dynamin relative to the start of the clathrin events, and lag time for the start of dynamin relative to the start of P/S. **c**. Relationship between clathrin assembly and membrane bending during CME.

Given that pol-TIRF could reliably detect nanoscale changes in membrane bending, we recorded the dynamics of membrane bending on single endocytic events in SK-MEL-2 cells labeled with DiI that endogenously express clathrin-Tq2 and dynamin2-eGFP (Extended Data Figure 5 and movies S1, S2). Single diffraction-limited endocytic events were tracked(21), filtered to retain only those that contained isolated clathrin, dynamin, and P/S events, and categorized based on detection of membrane bending relative to clathrin arrival (Fig. 3). The reliability of the detection of P/S relative to clathrin arrival can be seen in the example traces (Extended Data Figures 7-9). From these cells, approximately 7,100 tracks had clathrin-Tq2 signatures that appeared and disappeared during the time of imaging. Of these tracks, 481 were selected for analysis based on a set of criteria, the most stringent of which were the absence of adjacent membrane curvature in the P/S image and a dynamin signature (Extended Data Figure 6a). In approximately half of the CME events, membrane bending was detected at the moment clathrin arrived and grew in intensity (Fig. 3a, Class 1), consistent with bending during assembly. In the other half of the events, clathrin accumulated prior to detection of membrane bending (Fig. 3b, c). The delayed bending group was divided into two categories – a small subset in which all of the clathrin accumulated at the endocytic site prior to membrane bending (Fig. 3b, Class 2) and a larger group in which some clathrin accumulated prior to bending, but more clathrin was added during membrane bending (Fig. 3c, Class 3). The relative proportions of these events were Class 1 (43%), Class 2 (14%), and Class 3 (43%) (Fig. 3d). Thus, within the same cell, clathrin assembly and membrane bending occurs with heterogeneous timing with the clathrin coat accommodating several modes of membrane bending.

A population view of membrane bending dynamics during CME revealed a variable delay between clathrin assembly and the onset of membrane bending. We observed that the clathrin lifetimes for Class 1 events were shorter than Class 2 and 3 events (Class 1 (80±42 s), Class 2/3 (108±39 s), p<0.05, Fig. 4a). Unlike clathrin, the lifetimes of membrane bending and of dynamin showed no statistical differences across classes (Fig. 4a), indicating the dynamics of these processes are identical regardless of class. This suggested Class 2/3 events had a delay in progression that was not present in Class 1 events. Delays of unknown mechanism have been suggested for CME(22) and a checkpoint related to membrane bending has been proposed(21). In comparison with class 1, class2/3 events lagged behind the start of clathrin assembly with Δt =25.5 s (Fig. 4b). This same mean delay was also observed when comparing the lag for dynamin between class 1 and class 2/3 (Δt = 24.1s) (Fig. 4b).We observed minimal differences between the dyanimin lag from P/S initiation across the classes (Δt = 0.2 s, Fig. 4b), indicating that the principle difference in lifetime for the two classes arose during the time that clathrin began to assemble and the onset of membrane bending (Fig. 4c).

These results raise the possibility that the initial moments of clathrin association with the plasma membrane determine whether bending will begin immediately or if a flat intermediate state will occur. Correlative dSTORM-platinum replica TEM of clathrin structures containing only a few triskelia revealed both flat and curved morphologies (Fig. 1c), consistent with a bifurcation for entry into either Class 1 or Class 2/3 occurring early in the assembly process. Factors that influence this bifurcation could include the shape of the membrane at the moment clathrin binds, lateral membrane tension(23), curvature sensing/generating proteins(24), the stabilization of the curved state by additional factors such as AP2(25), or engagement of the actin cytoskeleton(22). Consistent with a stabilization step being required, we observed many short-lived (<18 sec) flat clathrin associations that did not recruit dynamin (Extended Data Figure 6b-d). Close inspection of the CME events did not reveal any examples of P/S signals that preceded clathrin association, suggesting that either initial membrane topography was not a factor in defining the sites at which clathrin assembled or that the scale of membrane bending needed to recruit clathrin was below what could be detected by pol-TIRF. Essentially, all CME structures that acquired P/S, acquired dynamin, indicating that once membrane bending starts, progression to a vesicle is a robust process (Fig. 4b). Thus, within the same cell, clathrin bends membrane through multiple heterogenous pathways in which the initiation of curvature is a key rate limiting step (Fig. 4c).

## Methods

### Reagents

1,1’-Dioctadecyl-3,3,3’,3’-tetramethylindocarbocyanine perchlorate (DiI; Sigma, St. Louis, MO) was dissolved in DMSO to prepare a 1 mg/mL stock. Cells were labeled using 1μg/mL DiI in 2.5% DMSO/dPBS. Imaging buffer was sterile 1x Dulbecco’s PBS with calcium and magnesium added (ThermoFisher, Rochester, NY) supplemented with 5mM glucose (Mediatech) and 10 mM Hepes (ThermoFisher). 100 μl DiI solution was added dropwise to 1 ml of imaging buffer and mixed by pipetting for less than 30 seconds and subsequently washed 3 times in imaging buffer and visualized by polarized TIRF immediately for no more than 30 minutes. Cells were imaged if p-polarized excitation intensities were less than 3000 to ensure that dynamics were not altered.

### Cell culture

Human SK-MEL-2 cells (Parental SK-MEL-2 and hDNM2^EN^) were a kind gift of D. Drubin (University of California, Berkeley, CA). Cells were cultured in DMEM (Hyclone, ThermoFisher) supplemented with 10% (v/v) fetal bovine serum (FBS, Hyclone), penicillin and streptomycin, and glucose. For live cell imaging experiments, cells were plated on fibronectin coated (Number 1.5) 25 mm coverslips (ThermoFisher) at approximately 50% confluency and imaged within 4-6 hours of plating. Flamed cleaned coverslips were coated with fibronectin at a final concentration of 25 μg/ml in dPBS for 30 minutes prior to plating cells. For visualization on the microscope, coverslips were transferred to AttoFluor chambers (ThermoFisher) and maintained in imaging buffer for up to 30 minutes.

### TIRF-based fluorescence microscopy

TIRF-based imaging was conducted using an inverted microscope built around a Till iMic (Till Photonics, Germany) equipped with a 60×1.49 N.A. oil immersion objective lens, diagrammed in Extended Data Figure 2. The microscope was enclosed in an environmental chamber and maintained at a temperature of 35-37°C using heater fans. Excitation for polarized TIRF was provided by a 561 nm laser. Excitation for mTurquoise2 and eGFP constructs was provided by 445 nm and 488 nm lasers, respectively. Lasers were combined into an acousto-optic-tunable filter and launched into a single mode fiber. Laser excitation was sent to a 2D scan head (Yanus, Till Photonics) which, along with a galvanometric mirror pair (PolyTrope, Till Photonics), was used to position the laser focal spot in the back focal plane of the objective lens for 2-point, and 360-TIRF illumination. A polychromic mirror reflected excitation wavelengths to the sample, zt442/514/561 was used for P/S and mTq2 excitation and zt405/488/561 was used for eGFP excitation (2.2 mm substrate, Chroma Technology, Bellows Falls, VT). Fluorescence emissions were first separated by a dichroic mirror (540/80dcrb, Chroma Technology) to send Green to one arm of the microscope where it was reflected to detector D3 (ZT514rcd, Chroma Technology) and Blue and Red fluorescence to the second arm, a second longpass dichroic (560 LPXR, Chroma Technology) allowed red fluorescence to pass to detector, D1, and reflected blue to detector, D2. Bandpass filters were used in front of each detector (D2 (Tq2) - ET470/24m, Chroma Technology, D3 (eGFP) – FF02-510/10m, Semrock, Inc, Rochester, NY, and D1 (DiI) – ET620/24m, Chroma Technology) and finally collected on three electron multiplying charge-coupled device cameras (iXon3 885, Andor Technology, Belfast, Ireland). The magnified pixel size is 133 nm a side. The exposure time was held constant at 100 ms for 442 and 488 excitations and 125 ms for P and S excitation. For 2-point images, the P and S images at each position were 62.5ms. Laser powers were measured using a PM120D digital handheld power meter (ThorLabs, Newton, NJ) and were typically between 0.5-20 mW during imaging. Bias calibration was performed by acquiring a set of 30 images with a closed shutter in front of each camera. The image stack was averaged to calculate the representative bias level in each pixel. Bias images were subtracted from each frame in the raw dataset.

### Back-focal plane centering

In order to determine the center of the objective lens’ optical axis, a calibration was carried out daily and for each chamber used in data acquisition. This centering calibration ensured that the excitation laser light encountered the glass/cell interface with a single incidence angle and, hence, produced a single TIRF excitation volume. A calibration protocol was used that steered the laser to positive and negative mirror positions, and thus incidence angles, with a small increment (1002 total steps), the intensity of the reflected light from the glass/water interface was read on a commercial quadrant photodiode module. Intensity values were plotted as a function of mirror position and the half-maximal values were used to adjust the mirror angles to center the optical axis (Extended Data Figure 2).

### Fiducial data collection and image registration

Images were registered using calibration images acquired simultaneously on each of the three EMCCD detectors. Briefly, 200 nm green beads (Life Technologies, Carlsbad, CA) immobilized on a glass coverslip were excited using 445 nm excitation and images were acquired as a single bead was moved across the field of view to create a well-sampled grid. Beads were localized in each channel and a polynomial transformation was determined to overlay each of the red and blue camera data onto that of the green detector. Because two excitation mirrors were used during the experiment fine tuning of the registration was completed using clathrin and dynamin spots that nearly correlate following the bead-based registration. Spots were identified, the inverse transformation was applied to obtain the original coordinates of the objects, and the subpixel positions were found by fitting to a 2D Gaussian. These points served as the fiducials for the final rigid affine transformation.

### Simulation

To predict the quantitative relationships between clathrin assembly and membrane bending signals from pol-TIRF, we created a discrete 3D simulation in MATLAB (The MathWorks Inc, Natick, MA) and DIPimage toolbox version 2.8 (Delft University of Technology, Delft, The Netherlands), building on our previous work for 3D microscopy simulations(19, 26, 27). The plasma membrane was represented as a plane that could either be bent into a sphere via a fixed radius of curvature (Fig. 1a, Extended Data Figure 1) or through progressive bending (Fig. 1b, Extended Data Figure 1), forming a vesicle of diameter = 100 nm.

For membrane bending during assembly, the forming clathrin pit was modeled as a sphere of fixed radius (r=50 nm) intersecting a plane. By shifting the center of the sphere along the z-direction we obtain the topographies outlined in Extended Data Figure 1a. In this case, clathrin is assumed to cover the spherical cap and ultimately the spherical vesicle. In the case of clathrin assembly preceding curvature, the pit is modeled as a circular patch of membrane emanating from the plane of plasma membrane. Here, the circular patch is designated to have uniform lateral clathrin intensity and the sphere is translated vertically over a progression of discrete radii. Thus, the vertical shift was set to z_shift_ = Area/2π/r_i_, where r_i_ ranged from 25,000 nm (slightly bent) to 50 nm (fully formed sphere). This produced the progression in outlined in Extended Data Figure 1b.

The fluorophores were modeled relative to each discrete element of plasma membrane or clathrin coat. Since 360 TIRF illumination was used for the clathrin images, no orientation dependencies were modeled. Thus, the equation defining the 2-dimensional clathrin image is given by,

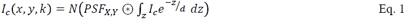

Where, I_c_ is the 3D (x,y,z) distribution of clathrin during pit stage k, d is the penetration depth of the TIRF field (100 nm), the microscope point spread function was modeled as a Gaussian distribution of with full width half max of 211 nm, typical of a 1.49 NA objective lens. Detection noise (N) was modeled by drawing intensities from a Poisson distribution.

Simulation of the pol-TIRF signals was achieved using the pol-TIRF fluorophore excitation equations of Axelrod and Anatharam(13) for the relative contributions of a plane and a sphere. Based on this work, we assume that the depth-dependent detection of emitted polarizations in the near field was approximately constant for a 1.49 NA objective and could therefore be neglected. Thus, using our discrete model, the polarization for a growing pit could be described as a plane and spherical components excited by either p-pol or s-pol illumination.

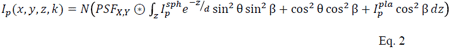

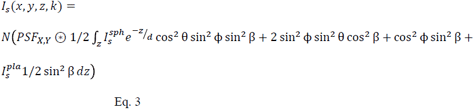

Where, spherical coordinates (θ,ϕ) are defined relative to the center of the sphere, β is the angle between the dipole moment of DiI and the plane, I^pla^ and I^sph^ are the intensities/unit membrane in the plane and sphere (assumed equal). Beta was determined by measuring planar regions of the plasma membrane and measuring the regional minimum which was found to be 0.26, which sets β=70°, which is nearly identical to the 69° value measured by Anantharam et al(13).

Discrete simulations were conducted on a 2 nm grid over pit morphological states, and then downsampled to nominal microscope pixel dimensions of 125 nm in x and y, and blurred with a Gaussian PSF with FWHM = 211 nm. Detection noise (N) was modeled by drawing intensities from a Poisson distribution. The code for this simulation will be made available via the Mathworks File Exchange.

### Correlative polTIRF-TEM and polTIRF-AFM

SK-MEL-2 expressing endogenous clathrin-Tq2 were plated on fibronectin coated coverslips for 5 hours, labeled with DiI, and sonicated with a Branson Digital Sonifier 450 in stabilization buffer (70mMKCl, 30 mM Hepes, 5mM MgCl2, at pH 4, and 1mM DTT) with a 1/8” tapered microtip approximately 5 mm above the coverslip for a single 400 ms pulse at 10% amplitude. Stabilization buffer was immediately removed after sonication using an aspirator and fixed with 2% formaldehyde, para (PFA) (Fisher Scientific) for 20 minutes followed by pol-TIRF imaging in PBS. In order to identify the same cells for correlative EM, a 9×9 grid of fluorescent images surrounding the cell of interest was collected, and a circle was drawn on the underside of the coverslip approximately around the imaged area using a high precision fine diamond scriber with a 0.5 mm diameter tip (Electron Microscopy Sciences). The coverslips were mounted on a slide with 10 μL of 2% glutaraldehyde to keep the sample hydrated (Sigma Aldrich). The coverslip was sealed with VALAP (1:1:1 mixture of Vaseline, lanolin, and paraffin) and epoxy prior to shipment for TEM or AFM.

### Electron Microscopy

Coverslips were transferred from glutaraldehyde into freshly prepared 0.1% w/v tannic acid in water and incubated at room temperature for 20 minutes. They were then rinsed 4x in water and transferred into 0.1% w/v uranyl acetate and incubated for 20 minutes, and rinsed with water prior to dehydration. Dehydration into ethanol, critical point drying, coating with platinum and carbon, replica lifting, and TEM were performed as previously described(28). Replicas were placed onto Formvar/carbon coated 75-mesh copper TEM grids (Ted Pella 01802-F).

### Atomic Force Microscopy

Correlated fluorescence-atomic force microscopy was performed using an Olympus IX73 inverted microscope equipped with an Olympus PlanApo 60X 1.45 N.A. oil immersion objective and Hamamatsu ORCA-Flash 4.0 V2 CMOS camera. The system was integrated with an Asylum Research MFP-3D-BIO AFM system and placed in a vibrational isolation chamber. AFM scanning of the unroofed cells in PBS solution was conducted at room temperature, under non-contact/AC mode, using Mikromasch CSC381/Cr-Au cantilevers, with a nominal spring constant of 50 pN/nm and a resonance frequency of 14 kHz.AFM image analysis was performed in Asylum Research AFM software. The correlative fluorescence-AFM images were aligned based on the correlation of spots between the two sets of clathrin fluorescence images to define fiducial markers for image registration, and quantified as described below.

### Image Correlation

Clathrin structures and positions were manual identified in the EM images by the appearance of the honeycomb lattice. The radius and centroid of each object was manually determined by fitting circles on the clathrin structures until it was completely encompassed. The ultrastructures were qualitatively categorized as [1] – Flat, [2] – Shallow curvature, [3] – Medium curvature, [4] – Domed curvature, [5] – Formed clathrin vesicle, based on the relative shadowing on the edge of the object. Finally, the coordinates of Tq2-Clathrin were obtained by fitting the spots to a 2D Gaussian, and identifying at least 4 spots to use as fiducial markers to generate the 2D affine transformation matrix that minimized the distance between correlated spots.

### Tilt-series tomography image analysis

The radius and heights of clathrin ultrastructures were quantified by manually fitting circles to the xy-sum projection to determine the radius, and drawing an arc along the yz-sum projection of the tomogram to determine the height.

### Correlative dSTORM - platinum replica transmission electron microscopy

SK-MEL-2 cells for correlative microscopy in Fig. 1 were obtained from ATCC and were grown in DMEM lacking phenol red (Life Technologies 31053-036) and supplemented 10% v/v FBS with 1% v/v Glutamax (Life Technologies 35050-061), 1% v/v Penicillin/Streptomycin (Invitrogen 15070-063), and 1% v/v sodium pyruvate (Sigma S8636). Cells were transfected with EGFP-clathrin light chain (a) using lipofectamine 2000 on day one. On day two, they were sorted to obtain only GFP containing cells and plated on coverslips embedded with gold nanoparticles (hestzig.com, part #600-200AuF). The coverslips had been coated with a 1:40 solution of fibronectin (Sigma F1141) in PBS for 30 minutes. On day three, the cells were unroofed and labeled as described below.

First, coverslips were rinsed in stabilization buffer (70 mM KCl, 30 mM HEPES brought to pH 7.4 with KOH, 5 mM MgCl2) for 2 minutes and unroofed by sonication in 2% paraformaldehyde (PFA) in stabilization buffer. They were then fixed in 2% PFA for 20 minutes. After rinsing with PBS, cells were placed in blocking buffer (3% bovine serum albumin in PBS) for one hour. They were immunolabeled with 11 nM Alexa Fluor 647 labeled GFP nanotrap (preparation described below) in blocking buffer for 45 minutes, rinsed in PBS, and post-fixed in 2% PFA for 20 minutes. Coverslips were then imaged in a sealed chamber containing blinking buffer (10% w/v glucose, 0.8 mg/mL glucose oxidase, 0.04 mg/mL catalase, 100mM 2-mercaptoethanol made fresh in PBS immediately before imaging). dSTORM was performed on a Nikon NSTORM system with 10 kW/cm^2^ 647 nm laser in TIRF illumination with 30,000 10 ms frames. A final image was created with Nikon Elements NSTORM analysis software with 5 nm pixel spacing.

After imaging, coverslips were marked with a diamond objective marker (Leica 11505059).The oil was cleaned off of the coverslip with 80 % ethanol. They were then stored in 2% glutaraldehyde in PBS and processed for EM the following day. EM processing and imaging was performed as described above. The gold nanoparticles that were embedded in the coverslips were visible in both dSTORM and EM and were therefore used as spatial fiducial markers. Three gold nanoparticles were used to map the fluorescence onto the EM image using an affine spatial transformation and nearest neighbor interpolation.

GFP nanotrap was expressed and purified as previously described(29). It was then labeled with Alexa Fluor 647 NHS ester (ThermoFisher 37573) using 2.4 molecules of dye for every one nanobody. These were purified using size exclusion chromatography and concentrated to 11 μM. SDS-PAGE indicated 1-4 dyes per nanobody. All GFP-nanotrap concentrations were estimated assuming an A_280_ extinction coefficient of 26930 M^-1^cm^-1^.

## Acknowledgments

This material is based on work supported by the National Science Foundation under CAREER Award to ADH, Grant No. 0953561, the National Science Foundation/EPSCoR Cooperative Agreement #IIA-1355423, the South Dakota Research and Innovation Center, BioSNTR, and by the State of South Dakota BOR CRGP to ADH and DMR-1337586 and DMR-1206908 to SS. Any opinions, findings, and conclusions or recommendations expressed in this material are those of the author(s) and do not necessarily reflect the views of the National Science Foundation. KAS and JWT were supported by the Intramural Research Program of the National Heart Lung and Blood Institute, National Institutes of Health in the Ack section. We thank the NHLBI, NIH electron microscopy and light microscopy core facilities for use of equipment. We thank Alan Hoofring of NIH Medical Arts for design work on Figures 1a,b and 2a

## Supplementary Materials

Movies S1-S2

